# Sex-and Stress-Dependent Plasticity of a Corticotropin Releasing Hormone / GABA Projection from the Basolateral Amygdala to Nucleus Accumbens that Mediates Reward Behaviors

**DOI:** 10.1101/2024.11.30.626183

**Authors:** Lara Taniguchi, Caitlin M Goodpaster, Gregory B de Carvalho, Matthew T Birnie, Yuncai Chen, Lulu Y Chen, Tallie Z Baram, Laura A DeNardo

**Author notes:** <u>Corresponding Authors:</u> Tallie Z Baram, Laura A DeNardo. equal contribution (Lara Taniguchi, Caitlin M Goodpaster). equal contribution (Tallie Z Baram, Laura A DeNardo).

## Abstract

**Background:** Motivated behaviors are executed by refined brain circuits. Early-life adversity (ELA) is a risk for human affective disorders involving dysregulated reward behaviors. In mice, ELA causes anhedonia-like behaviors in males and augmented reward motivation in females, indicating sex-dependent disruption of reward circuit operations. We recently identified a corticotropin-releasing hormone (CRH) expressing GABAergic projection from basolateral amygdala (BLA) to nucleus accumbens (NAc) that governs reward-seeking deficits in adult ELA males—but not females.

**Methods:** To probe the sex-specific role of this projection in reward behaviors, adult male and female CRH-Cre mice raised in control or ELA conditions received excitatory or inhibitory Cre-dependent DREADDs in BLA, and then clozapine N-oxide or vehicle to NAc medial shell during reward behaviors. We determined the cell identity of the projection using immunostaining and electrophysiology. Using tissue clearing, light sheet fluorescence microscopy and deep learning pipelines, we mapped brain-wide BLA CRH^+^ axonal projections to uncover sex differences in innervation.

**Results:** Chemogenetic manipulations in male mice demonstrated inhibitory effects of the CRH^+^ BLA-NAc projection on reward behaviors, whereas neither excitation nor inhibition influenced female behaviors. Molecular and electrophysiological cell-identities of the projection did not vary by sex. By contrast, comprehensive whole-brain mapping uncovered significant differences in NAc innervation patterns that were both sex and ELA-dependent, as well as selective changes of innervation of other brain regions.

**Conclusions:** The CRH/GABA BLA-NAc projection that influences reward behaviors in males differs structurally and functionally in females, uncovering potential mechanisms for the profound sex-specific impacts of ELA on reward behaviors.

## Introduction

Early-life adversity (ELA) arising from poverty, trauma, and chaotic environments affects millions of children worldwide (1). Studies link ELA to adverse cognitive and emotional outcomes (2–5), including disruptions in reward circuitry function (6–9) and impaired development of associated brain regions (10–12). Mental health disorders characterized by disruptions in the brain’s reward circuits often include anhedonia, a construct denoting decreased pleasure or desire for rewards (13). In humans, the relationship between ELA and dysregulated reward behaviors differs by sex (14,15), with women more susceptible to craving comfort food and developing eating and opioid use disorders (16,17) whereas men are more prone to alcohol use disorders (17,18). These differences may derive from sex-dependent characteristics of reward circuitry operations (14,15) which may be coupled with sex-biased developmental vulnerabilities to ELA (14,19–22). Establishing a direct causal link between ELA and anhedonia, and aberrantly augmented reward motivation in humans is challenging. Conversely, our rodent model of ELA consistently induces sex-specific disturbances in reward circuits, including anhedonia in males and augmented motivation for reward cues in females (8,23–27), providing a framework for addressing the mechanisms underlying these sex differences.

The brain is organized in circuits that execute complex behaviors, including those involved in reward. We hypothesized that circuits expressing stress-sensitive molecules such as the peptide corticotropin-releasing hormone (CRH), which is often involved in stress responses and adaptation (28–30), might be poised to be influenced by ELA. We initially mapped CRH projections to a key reward center in the brain, the nucleus accumbens (NAc), and identified a CRH-expressing (CRH^+^) GABA-ergic projection from the basolateral amygdala (BLA) to NAc shell (31). This was exciting because CRH modulates reward-related behaviors in NAc, and its actions there are context-dependent, influenced by prior or ongoing stress (32,33). We then established that activating the CRH^+^ BLA-NAc projection reduced reward behavior in adult control male mice, and silencing it in ‘anhedonic’ ELA male mice restored reward behaviors to control levels (27). This discovery is significant because it provides a mechanistic link between ELA and mood disorders (23,34).

Here, we investigate this novel, cell-type-specific projection’s role in modulating motivated reward behaviors in female mice and the fundamental sex- and ELA-dependent differences in the projection’s structure and function.

## Methods and Materials

### Mice

B6(Cg)-Crhtm1(cre)Zjh/J (CRH-ires-CRE, Jackson 012704) (35,36) mice were bred in-house, weaned and group-housed by sex on a 12-hour light-dark cycle (lights on 7 am). Post-surgery, single-housed males and group-housed females were moved to a reverse light-dark cycle (lights on 12 am). Procedures were approved by University of California-Irvine’s IACUC (AUP 24-104, 21-128) and adhered to NIH guidelines.

### Limited Bedding and Nesting ELA Model

To study ELA, we used the lab’s validated limited bedding and nesting (LBN) protocol (37,38,23). Dams and pups (4-8; sex balanced) were randomly assigned to cages with either customary or limited bedding and nesting materials. LBN cages had a plastic-coated mesh platform (McNichols 4700313244) ∼2.5 cm above the floor, and reduced bedding (150 ml), and nesting material (0.5 vs 1 nestlet). Cages were undisturbed during P2-P9 in a ventilated, quiet area. All mice returned to typical cages on P10 and weaned on P21.

### Surgeries

2-4 month old CRH-ires-Cre mice were anesthetized with 5% isoflurane and maintained at 1–1.5% on a stereotaxic machine with a heating pad. Crown fur was shaved, the skin sanitized (betadine and 70% ethanol), and skull exposed. Injection sites were drilled, and viral vectors were delivered via pulled pipettes (Drummond 2-000-010) at 100 nl/min, with pipet in place for 5 min. Post-surgery, mice received buprenorphine (Patterson Vet 07-894-9214) and were monitored for two days. To allow viral expression in axonal processes (31), we waited 6 weeks prior to chemogenetics and 8 weeks for optogenetics and brain clearing. For the latter, AAV1-EF1a-DIO-hChR2(H134R)-EYFP (Addgene 20298-AAV1) was injected into medial BLA bilaterally (0.2 µl/side; A/P −1.4mm, M/L ±4.25mm, D/V −4.60mm, 15° angle). For chemogenetic experiments, AAV2-hM3D(Gq)-mCherry (excitatory, Addgene 44361-AAV2) or AAV2-hM4D(Gi)-mCherry (inhibitory, Addgene 44362-AAV2) were bilaterally injected into medial BLA (0.2 µl/side). Guide cannulas were implanted bilaterally into medial NAc shell (A/P +1.2mm, M/L ±1.5mm, D/V −3.5mm, 12° angle). A dental cement headcap (Patterson Dental Orthojet Liquid 74594412; Powder 74598371) secured the cannulas with skull screws (Antrin AMS1201BINDSS).

### Behavioral and Chemogenetic Experiments

Behavior tests were performed within the first 2 hr of the active, dark phase (2,39) in a dimly lit room. All experiments were conducted in standard mouse cages (34 × 18 cm), either the home cage (males), or individual cages following habituation (females). For chemogenetic experiments, 0.2 µl clozapine-N-oxide (CNO, 1 mM, Hello Bio HB6149) (40,41) dissolved in saline (VetOne V1510223) was bilaterally infused into the medial NAc shell.

### Palatable Food Task

This assessed Cocoa Pebbles (Post) consumption in non food-restricted mice. Mice were habituated to Cocoa Pebbles for three days to minimize novelty effects (42) as follows: Day 1: ∼1 g Cocoa Pebbles placed in home cages overnight (consumption verified, not measured). Days 2–3: Mice placed in individual cages in the testing room for 1 hr, and 1 g Cocoa Pebbles provided. Consumption was measured after 1 hr. Test Days: Day 1: Mice received CNO or saline (counterbalanced) into the NAc, then moved to the behavior room where ∼1 g Cocoa Pebbles was provided, and a 1 hr consumption measured. Day 2: Same procedure, with counterbalanced CNO or saline. The one-day interval allowed CNO and vehicle washout (43), and counterbalancing controlled for infusion order effects.

### Sex Cue Preference

This task measured the active approach and interest of mice toward a Q-tip scented with urine (sex cue) versus one scented with almond (neutral cue). Female subjects were exposed to Q-tips scented with male urine (44), male subjects were exposed to Q-tips scented with estrous female urine (27). Urine was collected on the test day or stored, capped, at 4°C for <3 days. Duration of sniffing and frequency of approaches toward the urine- or almond-scented Q-tip were recorded. On Day 1, mice received CNO or saline (counterbalanced) infusion into the NAc, then moved to cages with Q-tips (60 µl each of urine and almond scents) placed at opposite corners. Exploration of the cues was assessed over 3 min. On day 2, the procedure was repeated with counterbalanced infusions (CNO or saline).

### Electrophysiology Slice Preparation

Mice were deeply anesthetized with isoflurane and quickly decapitated. Acute horizontal slices (300 µm) encompassing NAc and BLA were obtained using a vibratome (Leica V1200S) in an ice-cold sucrose cutting solution containing (in mM): 228 sucrose, 11 glucose, 26 NaHCO3, 1.2 NaH2PO4, 2.5 KCl, 5 Na-ascorbate, 3 Na-pyruvate, 10 MgSO4-7H2O, and 0.5 CaCl2 (305–310 mOsm, pH 7.4). Slices equilibrated in a homemade chamber for 25–30 min (34°C) then 45 min in room temperature aCSF containing (in mM): 119 NaCl, 26 NaHCO3, 1 NaH2PO4, 2.5 KCl, 11 glucose, 10 sucrose, 1.3 MgSO4-7H2O, and 2.5 CaCl2 (290–300 mOsm, pH 7.4), before being transferred to a recording chamber. Solutions were continuously bubbled with 95% O2/5%CO2.

### Whole-Cell Patch Clamp

Recordings were performed in the medial NAc shell using a Multiclamp 700B, Digidata 1550B, and Clampex 11 (Molecular Devices). Recordings were conducted in voltage-clamp mode at 31°C, low-pass filtered at 2 kHz, and digitized at 10 kHz. Borosilicate glass pipettes (3–4 MΩ, Molecular Devices) filled with internal solution (295–305 mOsm, pH 7.4, adjusted with CsOH) contained (in mM): 135 CsMeSO4, 8 CsCl, 10 HEPES, 0.25 EGTA, 5 Phosphocreatine, 4 MgATP, 0.3 NaGTP, and 1 mg/ml NeuroBiotin (295–305 mOsm, pH 7.4 with CsOH). Access resistance (Ra) was monitored, and recordings with Ra changes >20% were excluded. Cells were visualized with infrared DIC microscopy (Olympus BX51WI). Neurons were held at 0 mV to record optically evoked inhibitory postsynaptic currents (oIPSCs) using 488 nm LED light. After 5 minutes of stable oIPSCs, picrotoxin (100 µM) was superfused until oIPSCs were blocked. Neurons were then held at −60 mV to record optically evoked excitatory postsynaptic currents (oEPSCs). Stable oEPSCs were followed by the superfusion of CNQX (10 µM) and AP5 (50 µM). Slices were fixed in 4% PFA at 4°C overnight, then stored in 0.1M PBS for histological processing.

### Immunohistochemistry

Concurrent immunolabeling of CRH and GAD was performed as described previously (36,45). Briefly, sections were first incubated with a rabbit anti-CRH antiserum (PBL rC68, courtesy Dr. Paul Sawchenko, Salk Institute, La Jolla) (1:20,000, 7 days, 4°C), and then treated with HRP-conjugated anti-rabbit IgG (1:1,000, PerkinElmer) for 1.5 hours. Cyanine 3-conjugated tyramide (1:150, Akoya) was applied in the dark for 5-6 minutes. After CRH detection, sections were exposed to a mixture of mouse anti-GAD67 (1:250, Santa Cruz, sc-28376) and anti-GAD65 (1:1,000, Boehringer Mannheim, #1522 825) (3 days, 4°C), and visualized using anti-mouse IgG conjugated to Alexa Fluor 488 (1:400, Invitrogen).

### Brain Clearing

Brains were processed using the Adipo-Clear protocol (46) with slight modifications. Mice were transcardially perfused and brains hemisected ∼1 mm past the midline and postfixed overnight in 4% paraformaldehyde (Sigma Aldrich 30525-89-4) at 4°C. The following day, samples were dehydrated with a gradient of methanol (MeOH, Fisher Scientific 67-56-1)/B1n buffer (1:1000 Triton X-100, 2% w/v glycine, 1:10,000 NaOH 10N, 0.02% sodium azide) for 1 hr for each step (20%, 40%, 60%, 80%) on a nutator. Samples were then washed with 100% MeOH 2x for 1 hr each and then incubated in 2:1 dichloromethane (DCM):MeOH solution overnight. The following day, the samples were washed 2x for 1 hr in 100% DCM, followed by three washes of 100% MeOH (30 min, 45 min, 1 hr). Samples were bleached for 4 hr in 5:1 H2O2/MeOH buffer. A cascade of MeOH/B1n washes (80%, 60%, 40%, 20%; 30 min each) rehydrated the samples, followed by a 1 hr wash in B1n buffer. Tissue was permeabilized in 5% DMSO/0.3 M Glycine/PTxWH (1 hr then 2 hr). Samples were washed with PTxwH for 30 min and incubated in fresh PTxwH overnight. The following day, we performed two PTxwH washes (1 hr, then 2 hr). Samples were incubated in primary GFP antibody (GFP-1020, AVES Labs NC9510598) at 1:2000 in PtxwH shaking at 37°C for 11 days, washed in PtxwH (2x 1 hr, 2x 2 hr and then once per day for 2 days). Samples were then incubated in a secondary antibody (AlexFluor 647, ThermoFisher Scientific A78952) for 8 days shaken at 37°C. Samples were washed in PTxwH (same as after primary antibody). Samples were then washed in 1x PBS twice (1 hr, 2×2 hr, overnight). Samples were dehydrated in a gradient of MeOH:H_2_O (20%, 40%, 60%, and 80%; 30 min each and then 100% 30 min, 1 hr, 1.5 hr). Samples were incubated overnight in 2:1 DCM:MeOH on a nutator, washed in 100% DCM (2x 1 hr). Samples were cleared in 100% DBE. DBE was changed after 4 hr. Samples were stored in DBE in a dark location at room temperature. Imaging took place at least 24 hr after clearing.

### Whole-Brain Imaging

Brain samples were imaged on a light sheet microscope (Ultramicroscope II, LaVision Biotec) equipped with a sCMOS camera (Andor Neo) and a 2x/0.5 NA objective lens (MVPLAPO 2x) with a 6mm working distance dipping cap. Image stacks were acquired at 0.8x optical zoom using Imspector Microscope v285 controller software. We imaged using 488 nm (laser power 20%) and 640-nm (laser power 50%) lasers. The samples were scanned with a step-size of 3 µm using the continuous light-sheet scanning method with the included contrast adaptive algorithm for the 640-nm channel (20 acquisitions per plane), and without horizontal scanning for the 488-nm channel.

### DeepTraCE Pipeline

Whole-brain image stacks were analyzed using python based DeepTraCE GUI (https://github.com/jcouto/DeepTraCE). Briefly, image stacks in the 640-nm channel were segmented using the 3D U-net based machine learning pipeline TrailMap (47) with a model trained to recognize clearly delineated axons (Model 1) described in Gongwer et al., 2023. Following segmentation, the 488 nm autofluorescence channel and axon segmentation were converted to 8-bit, scaled to 10 µm resolution and rotated to match the reference atlas. The autofluorescence channel was registered using elastix to the Gubra Lab light sheet fluorescence microscopy (LSFM) atlas average template, which has annotations that match the Allen CCF (48–50). The same transformation was applied to the segmented brain using transformix and converted to 8-bit .tif format. Axons were then skeletonized (reduced to a single pixel thickness) for quantification, as described previously (47,51).

To account for variability in BLA viral expression across animals, we normalized all axon counts to the total number of labeled pixels across the whole brain (Fig. S1). Regional axon innervation was quantified by counting the number of skeletonized pixels in each brain region above a threshold, then dividing this pixel count by the total number of pixels in a region. Regions were defined by a collapsed version of the Gubra Lab LSFM atlas. The atlas was cropped on the anterior and posterior ends to match the amount of tissue visible in our data. Fiber tracts, ventricular systems, cerebellum, pons, medulla and olfactory bulb were excluded from analysis.

### Axon Quantification

Medial-lateral axon distributions within regions were calculated in MATLAB by binning the whole-brain image into 100 µm voxels, calculating the percentage of segmented pixels within each voxel normalizing for total fluorescence as above. The averaged summation of axon counts from each group was then averaged and plotted along with the SEM.

### Axon Visualization

To visualize axons in the NAc of representative brains (Fig. 3B), z-projections of raw light sheet were created in FIJI by scaling images to a 4.0625 µm space, virtually reslicing images in the coronal plane and performing maximum intensity z-projections of 30 µm depth followed by local contrast enhancement. To visualize axons at various coronal levels (Fig. 4A), representative brains from each group were resliced in the coronal plane, and then 80 µm thick z-projections of the skeletonized and registered axons were produced from the resliced brains.

### Sex Considerations

To consider the possible effects of the estrous cycle and associated hormonal fluctuations on the function of the projection and female reward behaviors, (52,53), females were swabbed for a vaginal smear (<5 seconds) during daily handling.

### Statistics

All statistical comparisons and analyses were performed in GraphPad Prism V10. The specific analyses can be found in the figure legends and Tables S1-3.

## Results

### ELA influences reward behaviors in a sex-dependent manner

As adults, male and female mice exposed to ELA exhibited profound sex-dependent changes in reward behaviors. Male ELA mice consumed less palatable food (Fig. 1B) and explored sex cues (scents of urine from a female mouse in estrous) less than control male mice (Fig. 1C). In contrast, adult female mice that were reared in ELA conditions showed augmented reward behaviors compared to control females, evidenced by increased consumption of Cocoa Pebbles (Fig. 1B) and increased number of approaches towards a sex cue (scent of urine from a male mouse) (Fig. 1D). Of note, in males there was a significant difference between ELA and controls in both the number of approaches to the sex cue (Fig. 1D) and the duration of sniffing (Fig. 1C). In females, by contrast, the ELA and control groups did not differ in the duration of sniffing (Fig. 1C), but diverged in the number of approaches (Fig. 1D), a measure of motivation for reward. This suggests a specific motivational deficit in ELA females leading to approaching the sex cue significantly less than controls.

**Fig. 1:**
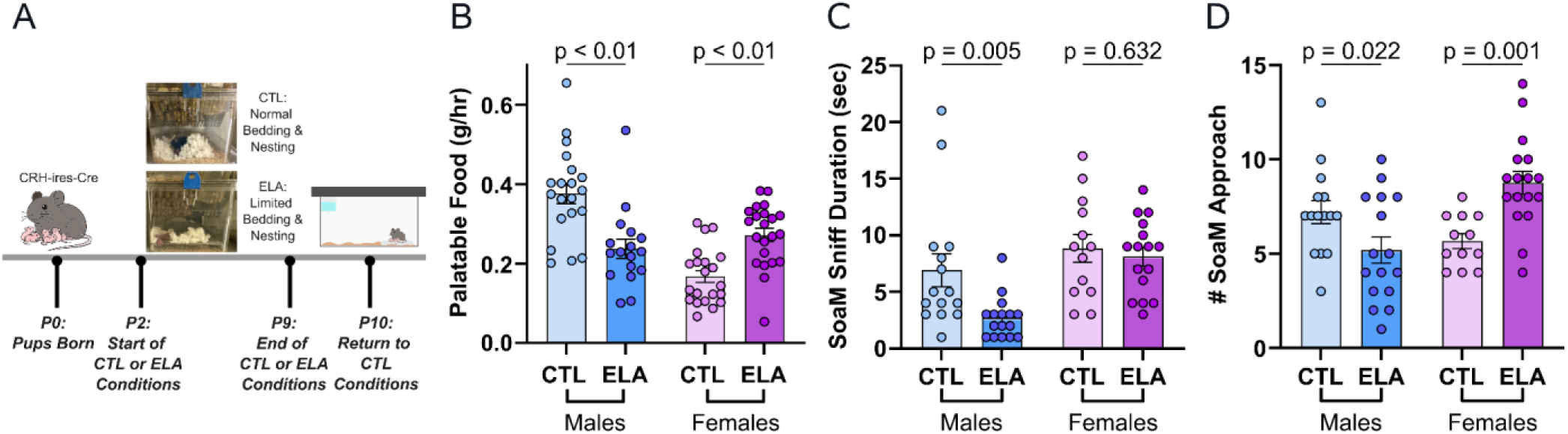
Sex differences in the impact of early life adversity (ELA) on adult reward behaviors. (**A**) Schematic of the limited bedding and nesting cages (LBN) model of ELA. Adult male mice raised in ELA cages exhibited “anhedonia-like” reward behavior compared to typically-reared mice (CTL) in (**B**) palatable food consumption, (**C**) duration spent sniffing a sex cue, and (**D**) number of approaches towards a sex cue. In contrast, adult ELA female mice had augmented motivated reward behaviors compared to control females, with increased (**B**) palatable food consumption and (**D**) number of approaches towards a sex cue, a measure of motivation, but not increased duration of sniffing, a measure of consummatory pleasure. In B-D, dots represent individual mice, error bars represent mean ± SEM. 2-way ANOVA. (**B**: n, CTL male = 20, CTL female = 21, ELA male = 17, ELA female = 22, **C**: n, CTL male = 15, CTL female = 13, ELA male = 16, ELA female = 16, **D**: n, CTL male = 15, CTL female = 12, ELA male = 16, ELA female = 17). CTL = control. ELA = early life adversity. LBN = limited bedding and nesting.

### The CRH expressing GABAergic BLA-NAc projection mediates reward behaviors in male but not female mice

In our recently published paper, we discovered the novel CRH^+^/GABA BLA-NAc pathway which inhibits reward-seeking behaviors and mediates the effects of ELA on reward behaviors in male mice (27). When the pathway was chemo- or optogenetically stimulated in control male mice, it suppressed reward behaviors. Inhibition of this projection in ‘anhedonic’ adult ELA male mice rescued reward behaviors. Here, we recapitulated these findings (Figs. 2B, 2C). Importantly, we report here the results of systematic manipulation of the projection in female mice, demonstrating profound sex differences. Chemogenetic activation of the CRH^+^ BLA-NAc pathway in control female mice did not affect palatable food consumption as observed in control males (Fig. 2B). Similarly, exciting the projection in ELA females had little effect (Fig. 2C). Chemogenetic <u>inhibition</u> of the CRH BLA-NAc pathway in female control and ELA mice failed to significantly alter palatable food consumption (Fig. 2D), in stark contrast to its effect in rescuing reward deficits in ELA males (27).

**Fig. 2:**
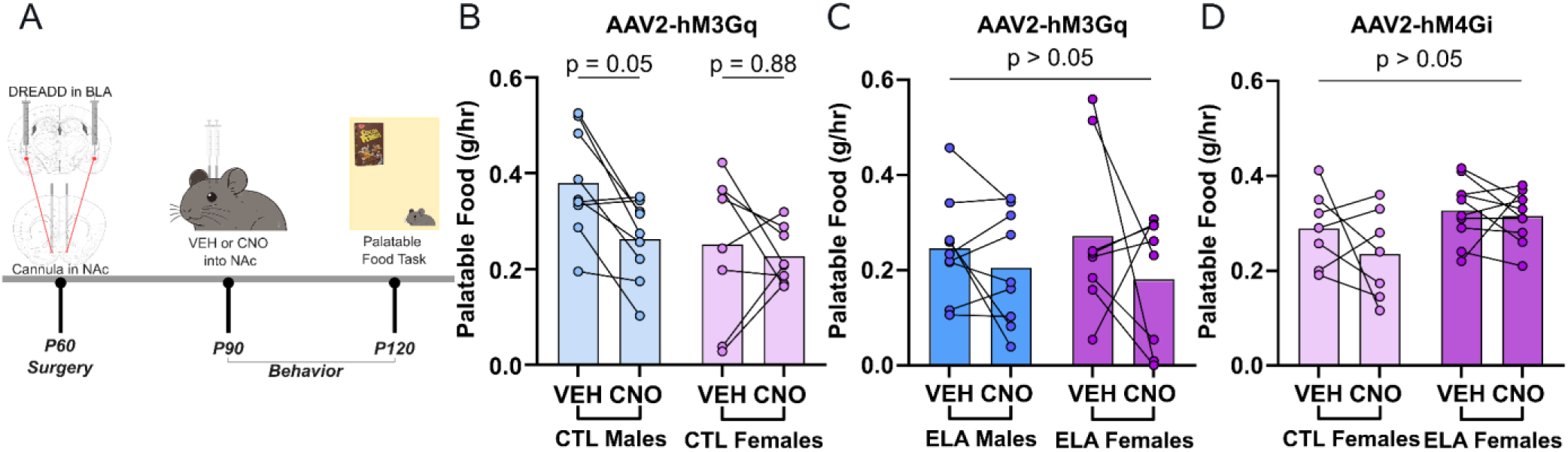
Chemogenetic manipulation of the CRH^+^ BLA-NAc projection *in vivo* influences reward behavior in male but not female control and ELA-exposed adult mice. (**A**) Viral strategy to chemogenetically target CRH^+^ BLA-NAc projection neurons. DREADD (Gq) excitation of CRH^+^ BLA-origin axon terminals in the NAc with 1 mM CNO reduced (**B**) palatable food consumption in control male mice but not in control female mice. (**C**) Excitation of the projection in ELA males, in which the pathway is overactive (27) has no effect, as is the case in female ELA mice. (**D**) Chemogenetic (Gi) inhibition of CRH^+^ BLA-origin axon terminals in the NAc with 1 mM CNO did not alter palatable food consumption in control or ELA female mice. In B-D, lines represent individual mice, bars represent mean. 2-way RM-ANOVA (**B**: *n,* CTL male = 9, CTL female = 8; **C**: *n,* ELA male = 9, ELA female = 8; **D**: *n,* CTL female = 7, ELA female = 10). VEH = vehicle (saline). CNO = clozapine N-oxide. CTL = control. ELA = early life adversity.

### The CRH^+^ BLA-NAc projection is GABAergic in both male and female mice

In male mice, CRH^+^ BLA-NAc neurons made GABA-ergic synapses onto target NAc cells and inhibited reward behaviors. In females, activating the projection had no detectable effects on reward behaviors. Therefore, we tested if the cell-identity of the projection might differ in females vs males. We injected CRH-ires-Cre mice with DIO-ChR2 in the BLA, and obtained whole-cell patch-clamp recordings from target neurons in the NAc medial shell. Inhibitory postsynaptic currents (IPSC) were optically-evoked with a 2 ms light pulse in the NAc (Fig. 3). Examples of the representative IPSCs traces are shown in Figs. 3A and 3E. After 5 min, the GABA^A^ receptor antagonist picrotoxin was washed on, blocking IPSCs in both males and female slices across all groups as apparent in the time-course data of normalized oIPSCs amplitudes (Figs. 3B, 3F) and time-point analysis (Figs. 3C, 3G). Raw oIPSCs amplitudes pre and post picrotoxin are shown in Figs 3D and 3H. These data indicate that in both male and female mice, optically evoked currents from CRH^+^ BLA-origin axons in the NAc are inhibitory and are mediated by GABA^A^ receptors. In further support of the GABAergic nature of the CRH^+^ BLA projection, all CRH cells in the BLA expressed the GABAergic markers GAD 65/67 (Figs. 3I-K).

**Fig 3.**
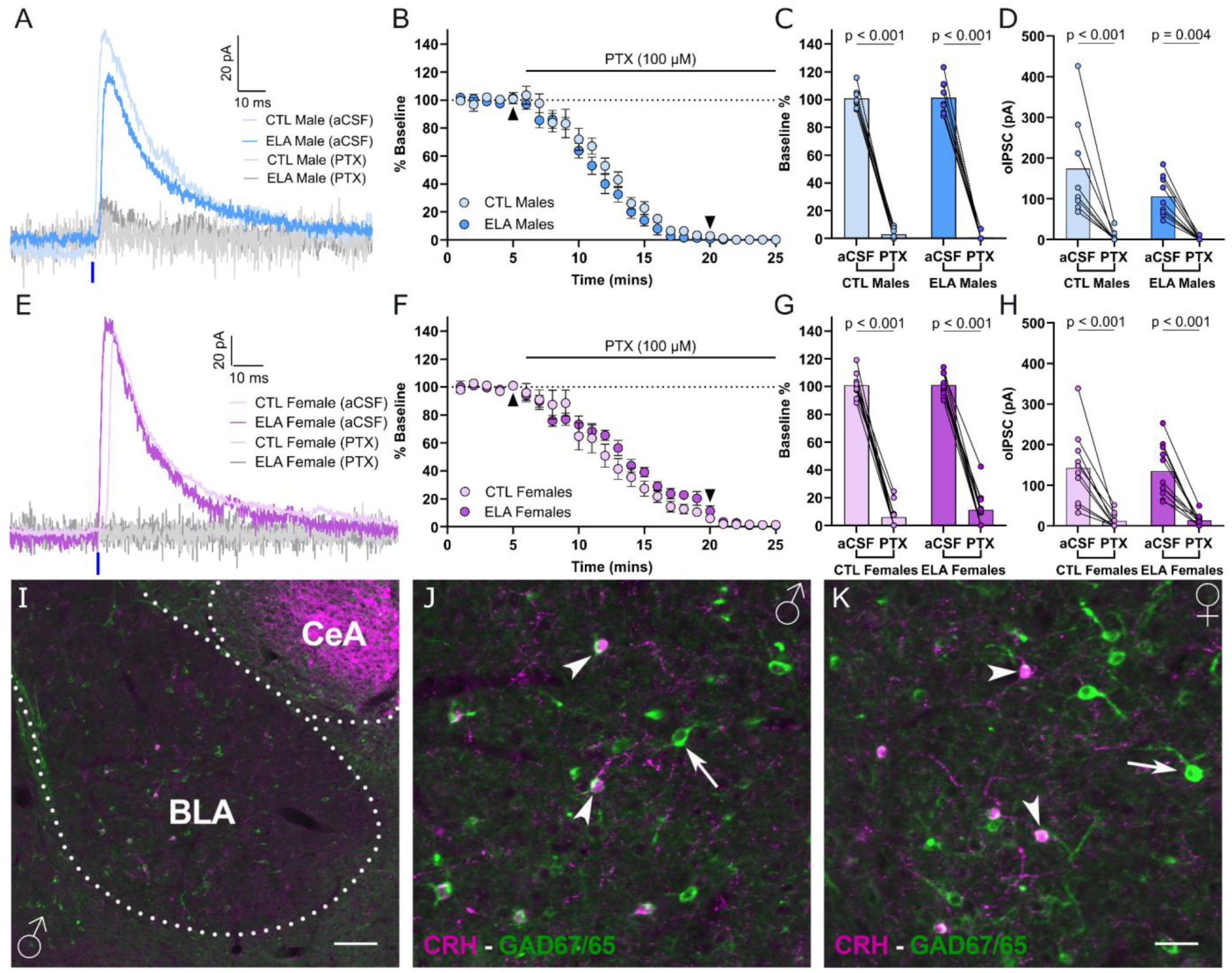
CRH^+^ BLA-NAc neurons are GABAergic, apparent from both electrophysiological and neurochemical markers. Whole-cell patch-clamp recordings were obtained from NAc medial shell neurons while inhibitory postsynaptic currents (IPSCs) were optically-evoked from CRH^+^ BLA neurons with a 2 ms light pulse at 488 nm. Representative traces of optically evoked IPSCs (oIPSCs) in males (**A**) and females (**E**). Time-course plot of normalized oIPSCs amplitudes before and after application of picrotoxin in (**B**) males and (**F**) females. Timepoint analysis of normalized oIPSC amplitudes at 5 min after start of recording compared to 15 min after picrotoxin application for (**C**) males and (**G**) females. (**D**,**H**) oIPSCs amplitudes pre and post picrotoxin. In B and F dots and error bars represent mean ± SEM. In C-D and G-H, dots represent individual cells, bars represent mean. 2-way RM-ANOVA (**B-D**: n, CTL male = 8 cells/4 mice, ELA male = 9 cells/4 mice; **F-H**: n, CTL female = 10 cells/4 mice; ELA female = 11 cells/4 mice). (**I-K**) Concurrent immunolabeling of CRH and both isoforms of GAD (67 and 65) in the BLA of a CTL male (**J**) and CTL female (**K**). High magnification photomicrographs indicate that all CRH expressing cells (magenta) also express GAD (arrowheads), whereas numerous GABAergic cells are devoid of CRH expression (arrows). In I, scale bars = 100 µm. In J-K, scale bars = 35 µm. aCSF = artificial cerebrospinal fluid. PTX = picrotoxin. CTL = control. ELA = early life adversity.

### Sex and rearing-dependent axonal innervation patterns of BLA origin CRH^+^ cells within the NAc

The results above indicate that there are sex-dependent differences in the function of the CRH^+^ BLA-NAc projection that are not explained by neurotransmitter identity or function of the originating cells. To investigate if structural differences in this pathway underlie these differences, we utilized a whole brain imaging approach to determine if sex or rearing alters the innervation of CRH^+^ BLA axonal projections within the NAc. Using a viral approach, we fluorescently labeled CRH^+^ BLA axons in adult male and female CRH-cre mice who experienced control or ELA rearing (Fig. 4A). We then cleared the brains, imaged them using light sheet fluorescence microscopy and quantified the brain-wide axonal projections of CRH^+^ BLA neurons (Fig. 4B).

**Fig. 4:**
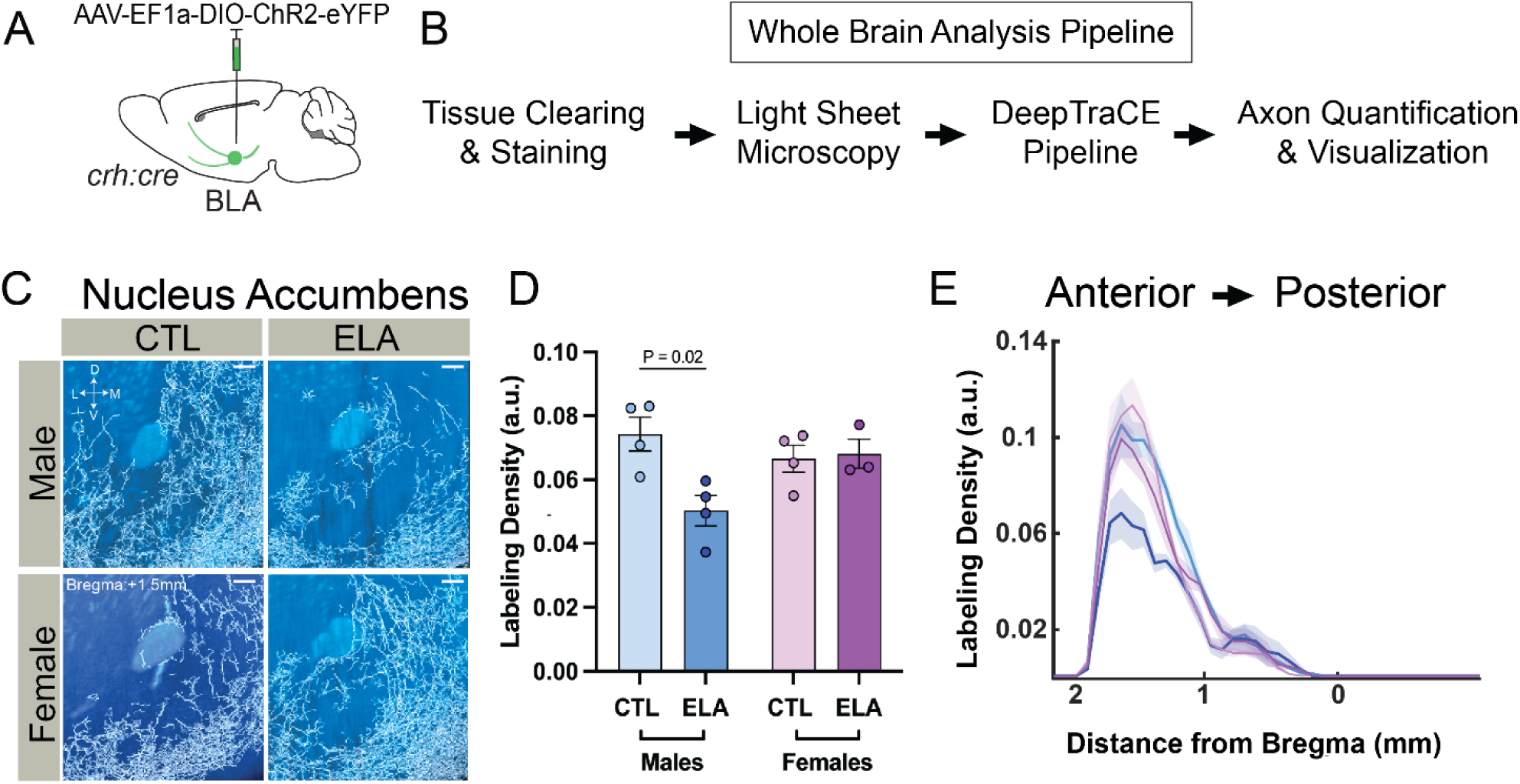
Axonal innervation patterns of CRH^+^ BLA cells within the nucleus accumbens (NAc). (**A**) Viral strategy for labeling axonal projections of CRH^+^ BLA neurons and (**B**) whole brain analysis pipeline. (**C**) Coronal view of 80 µm z-projections of skeletonized axons determined by DeepTraCE (axons, white) from example brains. In C, scale bars = 200 µm. (**D**) Differences in sex and rearing specific innervation patterns in NAc and quantification of differences in axonal innervation patterns among all groups. In D, error bars represent mean ± SEM. 2-way ANOVA (**D**: n, CTL male = 4, CTL female = 4, ELA male = 4, ELA female = 3). (**E**) Quantification of axonal innervation density along the anterior-posterior axis of the NAc. CTL = control. ELA = early life adversity.

Given the surprising differences in the functional role of the CRH^+^/GABA BLA-NAc pathway in reward behavior, we first focused our analysis on the NAc. In this region, both sex and rearing influenced the density of CRH^+^ BLA axonal innervation (Figs. 4C-E). Overall, ELA reduced labeling of axons in the NAc of male mice, with no effect on females (Figs. 4C-D). Notably, when comparing innervation of the NAc along the anterior-posterior axis, we found that this reduction in ELA males was most notable in the anterior NAc, with negligible changes in the posterior NAc (Fig. 4E). These changes were not a result of differences in the numbers of CRH^+^ neurons in the BLA of the four groups studied (Fig. S1), and are in line with the sex-dependent functional role of the CRH^+^ BLA-NAc pathway in motivated behavior.

### Sex and rearing impact brain-wide projection patterns of BLA CRH^+^ cells across the whole brain

Our results above, showing a rearing effect in males but not in females, on axonal innervation of CRH^+^ BLA neurons within the NAc correlate with, and potentially contribute to the reduction in reward behaviors in males who experience ELA compared to standard reared males. As no differences in axon labeling was observed in the NAc of female mice, we speculated whether ELA might induce anatomical changes in CRH^+^ BLA neuronal innervation in other downstream regions. To investigate this, we expanded our analysis to compare brain-wide projection patterns of CRH^+^ neurons in the BLA in typically-reared and ELA mice.

The axonal projection patterns of CRH^+^ BLA neurons were consistent with previous reports of BLA connectivity (Figs. 5A-B). For example, we found that these neurons project to the temporal association, the hippocampal formation, including the entorhinal cortex, CA1, and CA3 (54–56), the nucleus of the solitary tract, the piriform cortex, and the zona incerta (57,58). Unlike the overall BLA cell population, CRH^+^ cells also robustly innervated hypothalamic areas, including the paraventricular nucleus, anterior hypothalamic nucleus, ventral mammillary nucleus and the medial preoptic area.

**Fig. 5:**
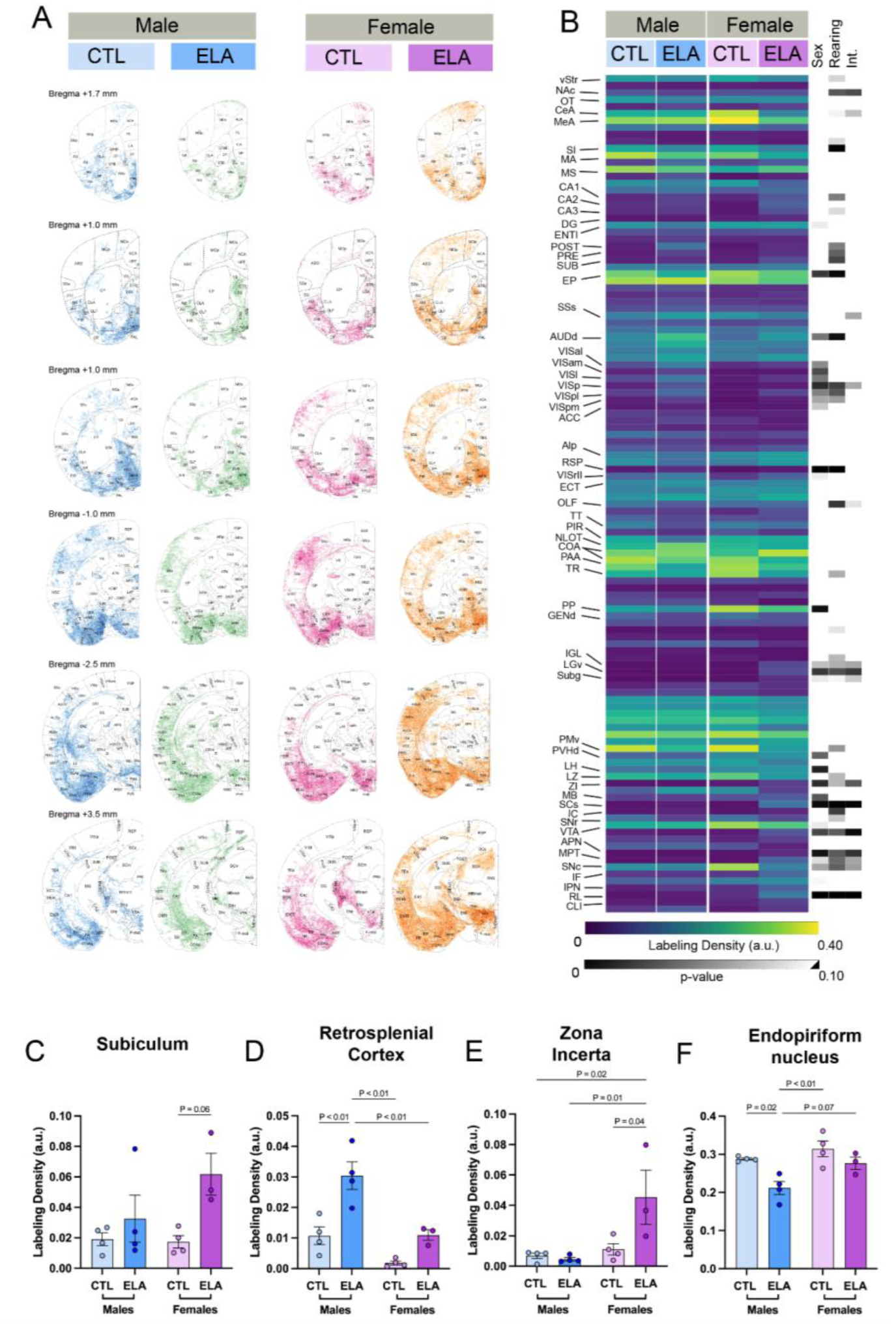
Visualization and quantification of brain-wide projection patterns of CRH^+^ BLA cells. (**A**) 80 µm coronal optical sections showing CRH^+^ BLA axons registered to the standardized brain atlas. Each image represents one example brain from each group. Innervation of BLA CRH^+^ cells in CTL males, ELA males, CTL females and ELA females are shown in blue, green, pink and orange respectively. (**B**) Left heatmap, relative labeling density averaged across conditions (normalized to region volume and total labeling) of 120 regions defined by the Allen Brain Atlas. Right heatmap, p values from 2-way ANOVA of axon innervation density comparing impact of sex, rearing and their interaction in each individual brain region. Sex and rearing effect on labeling density of axons in the (**C**) subiculum, (**D**) retrosplenial cortex, (**E**) zona incerta and (**F**) endopiriform cortex. 2-way ANOVA (B-F: n, CTL male = 4, CTL female = 4, ELA male = 4, ELA female = 3). CTL = control. ELA = early life adversity.

We next performed quantitative comparisons of axonal innervation density in each individual brain region, and identified several regions in which CRH^+^ BLA axonal density was selectively altered in females that experienced ELA. Salient examples include the subiculum, retrosplenial cortex and zona incerta–which are involved in fear, motivation and memory (59–61). Within the subiculum, ELA females display greater axonal innervation than control females and both male groups (Fig 5C). In the retrosplenial cortex, although CRH^+^ BLA axonal innervation density was relatively low, there was significantly more labeling in males who experienced ELA compared to control males and both female groups (Fig 5D). Additionally, the zona incerta labeling was significantly greater in females who experienced ELA compared to all other groups (Fig 5E). In contrast, reduced labeling in ELA males was found in the endopiriform nucleus, a region that is extensively connected with the limbic system (62) (Fig 5F).

## Discussion

The principal findings of the current studies are: (1) Unlike in males, chemogenetic manipulations of the CRH/GABA BLA-NAc projection in females do not influence reward behaviors. (2) Optical stimulation and immunohistochemistry indicate that the basic cell types and synaptic functions of the projection are not sex-specific. (3) The structural organization of the CRH^+^/GABA BLA-NAc projection is sex-specific and is further modulated by early-life rearing conditions. (4) Beyond projections to the NAc, the innervation patterns of CRH^+^ BLA neurons suggest global sex-dependent differences in their brain-wide organization, which might account for both fundamental differences in their operations as well as sex-dependent differences in the outcomes of ELA.

In male mice, we previously discovered that the CRH^+^/GABA BLA-NAc projection mediates the anhedonic-like effects of ELA on reward behaviors (27). Here we recapitulated these results, finding that chemogenetic excitation of the projection in typically reared adult male mice reduced palatable food consumption. In contrast, chemogenetic excitation of the projection did not affect palatable food consumption in either control or ELA females. While inhibiting the projection in ELA males rescued reward deficits (27), inhibiting the projection in either typically reared or ELA-reared adult females failed to influence palatable food consumption. These surprising results indicate profound, sex-specific differences in the functional roles of the CRH^+^/GABA BLA-NAc projection in male and female mice.

Reward behaviors include motivation (‘wanting’) and consummatory (‘liking’) aspects (63). Here we distinguished the motivational vs consummatory components of these reward behaviors in a second reward seeking behavior, the response to sex-reward cues. ELA reduced reward behaviors towards the scent of an estrous female in male mice and enhanced these behaviors in ELA females presented with the scent of a male mouse. Unexpectedly, we found sex-specificity in the effects of ELA on discrete components of these behaviors: ELA reduced consummatory behaviors (sniff times) and motivation (approaches to the scent-bearing Q-tip) in males. In contrast, in females, motivation for the sex-reward cue was augmented, with little change in sniff durations (Fig. 1). Because motivational and consummatory aspects of reward behaviors have distinct anatomical substrates, these findings suggest a need for investigating the nuanced connectivity and potential sex differences in the innervation and roles of the CRH^+^/GABA BLA-NAc projection.

We considered several variables that might have influenced our results. First, we excluded the possibility that the difference in signal density might be a result of differences in the number of CRH^+^ cells in the BLA of ELA and control female mice (Fig. S1). In addition, the estrous cycle stage of the experimental mice had no influence on the data and there was no litter effect. We also verified cannulae placements and injection sites, excluding mice with misplaced injections (Fig. S2). We then reasoned that the sex-dependent difference in the behavioral functions of the projection might be a result of neuroanatomical differences or neurotransmitter identity of the CRH^+^ BLA cells, and this hypothesis was refuted using immunohistochemistry (Fig. 3). The exclusion of such explanations for the sex-specific behavioral effects of ELA brought up the possibility that these functional differences derive from sex-dimorphic differences in either the functional properties or the structural organization of the projection.

We did not find sex- or rearing effects on the basic functional properties of the CRH/GABA cells projecting to the NAc. Using optogenetics paired with electrophysiology, we found that activating CRH cells in the BLA triggered inhibitory currents in the medial shell of the NAc across all groups, regardless of sex or rearing conditions (Fig. 3). All CRH^+^ BLA cells were GABAergic, indicating that the fundamental cell type and function of the projection are the same across sexes and rearing. Future studies should untangle the specific interactions between GABA and CRH in this projection and determine whether GABA, CRH, or both mediate the effects of the CRH^+^/GABA BLA-NAc projection on behaviors in both typically reared and ELA-reared mice of both sexes.

As inhibition of the CRH^+^/GABA BLA-NAc projection restored typical reward responses to a palatable food cue, we hypothesized that changes in this circuit may be due to structural remodeling that occurs following ELA. Specifically we reasoned that CRH/GABA BLA neurons may have increased NAc axonal innervation, thus leading to increased synaptic activity that influences NAc operations. To our surprise, we observed a decrease in axon labeling in the NAc of male mice. One possible explanation for this is that stress during early periods of development induces overactivation of this cell type with increased release of CRH, GABA or both. In turn, the brain undergoes structural plasticity to try to reduce activity and maintain homeostasis by reducing the strength or the number of synaptic connections in this pathway. Future studies are needed to examine this potential mechanism.

Beyond the NAc, our brain-wide analysis suggests that there is extensive target- and sex-specificity in the effects of ELA on BLA innervation patterns. For instance, in the subiculum and zona incerta, we observed female-specific increases in CRH^+^ BLA axons following ELA. In contrast, we observed male-specific increases in axon labeling in the retrosplenial cortex and male-specific decreases in axon labeling in the endopiriform nucleus. This suggests that subiculum and zona incerta-dependent behaviors may be more affected in females that experienced ELA whereas retrosplenial- and endopiriform-dependent behaviors may be more affected in males that experienced ELA. The subiculum is the output center of the hippocampus and is implicated in spatial processing, motivation, stress integration and anxiety-like behaviors (64,65). Previous studies of ELA reported a reduction in subiculum volume in humans and an early decline in neuronal growth markers in mice, but these studies did not examine sex differences (66,67). The zona incerta, an extension of the reticular formation of the thalamus, is a largely inhibitory nucleus with a heterogeneous cell population expressing a variety of different molecular markers (58). Recent studies have found the zona incerta to be important for integrating multisensory inputs, controlling motor behavior and driving motivational and appetitive behaviors (68,69). Altered zona incerta activity contributes to anxiety-related and coping behaviors following chronic stress (70,71), but the effects of ELA on zona incerta functions are poorly understood. The retrosplenial cortex plays a key role in spatial and contextual learning and memory (72,73), but little is known about how its function may be impacted by ELA. The endopiriform nucleus is extensively interconnected with the limbic system (74). While its functions are poorly understood, it may augment limbic circuit excitability (62) and may contribute to recognition memory (75). Overall, the effects of ELA on all four of these regions are poorly understood, so our anatomical findings warrant further functional studies.

Our observations of region-specific increases in axon labeling are consistent with previous studies that observed elevated BLA axon density in the prefrontal cortex of adolescent rats that experienced ELA (76). Several mechanisms may contribute to the increased axon labeling we observed. The increased density may result from augmented axon sprouting or a failure of developmental pruning mechanisms. The latter mechanism is supported by our prior work on the failure of synaptic pruning onto CRH^+^ cells in the hypothalamic paraventricular nucleus (77). The deficient synaptic pruning was a result of ELA-induced microglial dysfunction. By contrast, in the hippocampus, ELA has led to selective deficits of neuronal arborization in specific hippocampal layers (78,79) thus, ELA-induced dysregulation of structural brain-circuit maturation is a plausible common theme for the altered innervation observed here, and for the behavioral consequences (80).

## Acknowledgements and Disclosures

This work was supported by National Institute of Health grants: RO1 MH 132680 and P50 MH096889. We would like to thank Amanda Chiang and Anamika Paul for their technical assistance. All authors report no biomedical financial interests or potential conflicts of interest.

---

**Supplemental Table 1:** ANOVA Tables.

| Figure | ANOVA Table | SS (Type III) | DF | MS | F (DFn, DFd) | P value |
| --- | --- | --- | --- | --- | --- | --- |
| **Figure 1B** | *Interaction* | 0.2947 | 1 | 0.2947 | F (1, 76) = 35.21 | P<0.01 |
|  | *Sex* | 0.1518 | 1 | 0.1518 | F (1, 76) = 18.14 | P<0.01 |
|  | *Rearing* | 0.005957 | 1 | 0.005957 | F (1, 76) = 0.7117 | P=0.40 |
| **Figure 1C** | *Interaction* | 46.3 | 1 | 46.3 | F (1, 56) = 2.88 | P=0.095 |
|  | *Sex* | 201 | 1 | 201 | F (1, 56) = 12.5 | P<0.001 |
|  | *Rearing* | 91.9 | 1 | 91.9 | F (1, 56) = 5.72 | P=0.020 |
| **Figure 1D** | *Interaction* | 96.3 | 1 | 96.3 | F (1, 56) = 17.1 | P<0.001 |
|  | *Sex* | 15.4 | 1 | 15.4 | F (1, 56) = 2.74 | P=0.103 |
|  | *Rearing* | 4.34 | 1 | 4.34 | F (1, 56) = 0.773 | P=0.383 |
| **Figure 2B** | *Interaction* | 0.01845 | 1 | 0.01845 | F (1, 15) = 2.463 | P=0.14 |
|  | *Sex* | 0.05684 | 1 | 0.05684 | F (1, 15) = 3.739 | P=0.07 |
|  | *CNO* | 0.0423 | 1 | 0.0423 | F (1, 15) = 5.648 | P=0.03 |
| **Figure 2C** | *Interaction* | 0.005163 | 1 | 0.005163 | F (1, 15) = 0.2957 | P=0.59 |
|  | *Sex* | 0.00001398 | 1 | 0.00001398 | F (1, 15) = 0.0007340 | P=0.98 |
|  | *CNO* | 0.03655 | 1 | 0.03655 | F (1, 15) = 2.093 | P=0.17 |
| **Figure 2D** | *Interaction* | 0.003548 | 1 | 0.003548 | F (1, 15) = 0.7343 | P=0.40 |
|  | *Rearing* | 0.0289 | 1 | 0.0289 | F (1, 15) = 5.275 | P=0.04 |
|  | *CNO* | 0.008945 | 1 | 0.008945 | F (1, 15) = 1.851 | P=0.19 |
| **Figure 3C** | *Interaction* | 15.7 | 1 | 15.7 | F (1, 15) = 0.266 | P=0.614 |
|  | *Rearing* | 5.52 | 1 | 5.52 | F (1, 15) = 0.107 | P=0.748 |
|  | *PTX* | 83445 | 1 | 83445 | F (1, 15) = 1408 | P<0.001 |
| **Figure 3G** | *Interaction* | 68.7 | 1 | 68.7 | F (1, 19) = 0.721 | P=0.406 |
| | *Rearing* | 72.1 | 1 | 72.1 | $F(1, 19) = 0.758$ | $P=0.395$ |
| | *PTX* | 89380 | 1 | 89380 | $F(1, 19) = 939$ | $P<0.001$ |
| **Figure 3E** | *Interaction* | 8158 | 1 | 8158 | $F(1, 15) = 2.25$ | $P=0.154$ |
| | *Rearing* | 11714 | 1 | 11714 | $F(1, 15) = 2.31$ | $P=0.149$ |
| | *PTX* | 156529 | 1 | 156529 | $F(1, 15) = 43.1$ | $P<0.001$ |
| **Figure 3H** | *Interaction* | 215 | 1 | 215 | $F(1, 19) = 0.0844$ | $P=0.775$ |
| | *Rearing* | 127 | 1 | 127 | $F(1, 19) = 0.0328$ | $P=0.858$ |
| | *PTX* | 165342 | 1 | 165342 | $F(1, 19) = 65.0$ | $P<0.001$ |
| **Figure 4D** | *Interaction* | 0.0006027 | 1 | 0.0006027 | $F(1, 11) = 6.942$ | $P=0.02$ |
| | *Sex* | 0.00009448 | 1 | 0.00009448 | $F(1, 11) = 1.088$ | $P=0.32$ |
| | *Rearing* | 0.0004658 | 1 | 0.0004658 | $F(1, 11) = 5.365$ | $P=0.04$ |
| **Figure 5C** | *Interaction* | 0.0008831 | 1 | 0.0008831 | $F(1, 11) = 2.203$ | $P=0.17$ |
| | *Sex* | 0.0006967 | 1 | 0.0006967 | $F(1, 11) = 1.738$ | $P=0.21$ |
| | *Rearing* | 0.003087 | 1 | 0.003087 | $F(1, 11) = 7.699$ | $P=0.02$ |
| **Figure 5D** | *Interaction* | 0.0001021 | 1 | 0.0001021 | $F(1, 11) = 3.022$ | $P=0.11$ |
| | *Sex* | 0.0007418 | 1 | 0.0007418 | $F(1, 11) = 21.97$ | $P<0.01$ |
| | *Rearing* | 0.0007684 | 1 | 0.0007684 | $F(1, 11) = 22.76$ | $P<0.01$ |
| **Figure 5E** | *Interaction* | 0.001226 | 1 | 0.001226 | $F(1, 11) = 6.324$ | $P=0.03$ |
| | *Sex* | 0.001875 | 1 | 0.001875 | $F(1, 11) = 9.672$ | $P<0.01$ |
| | *Rearing* | 0.0009432 | 1 | 0.0009432 | $F(1, 11) = 4.866$ | $P=0.05$ |
| **Figure 5F** | *Interaction* | 0.001408 | 1 | 0.001408 | $F(1, 11) = 1.517$ | $P=0.24$ |
| | *Sex* | 0.007741 | 1 | 0.007741 | $F(1, 11) = 8.338$ | $P=0.01$ |
| | *Rearing* | 0.01185 | 1 | 0.01185 | $F(1, 11) = 12.76$ | $P<0.01$ |
| **Figure S1B** | *Interaction* | 0.0005008 | 1 | 0.0005008 | $F(1, 11) = 1.116$ | $P=0.31$ |
| | *Sex* | 0.0000661 | 1 | 0.0000661 | $F(1, 11) = 0.1473$ | $P=0.71$ |
| | *Rearing* | 0.003156 | 1 | 0.003156 | $F(1, 11) = 7.033$ | $P=0.02$ |

**Supplemental Table 2:**
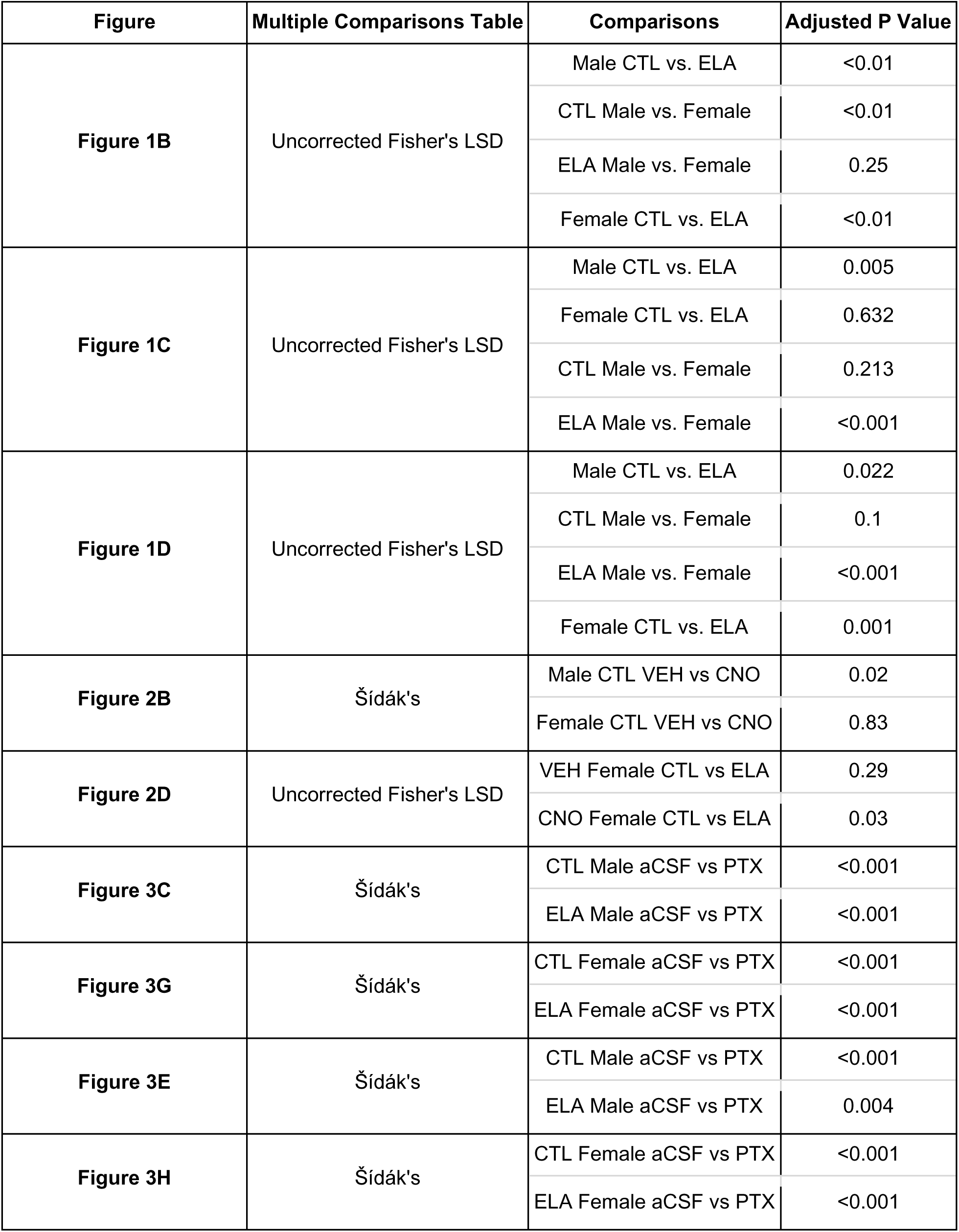
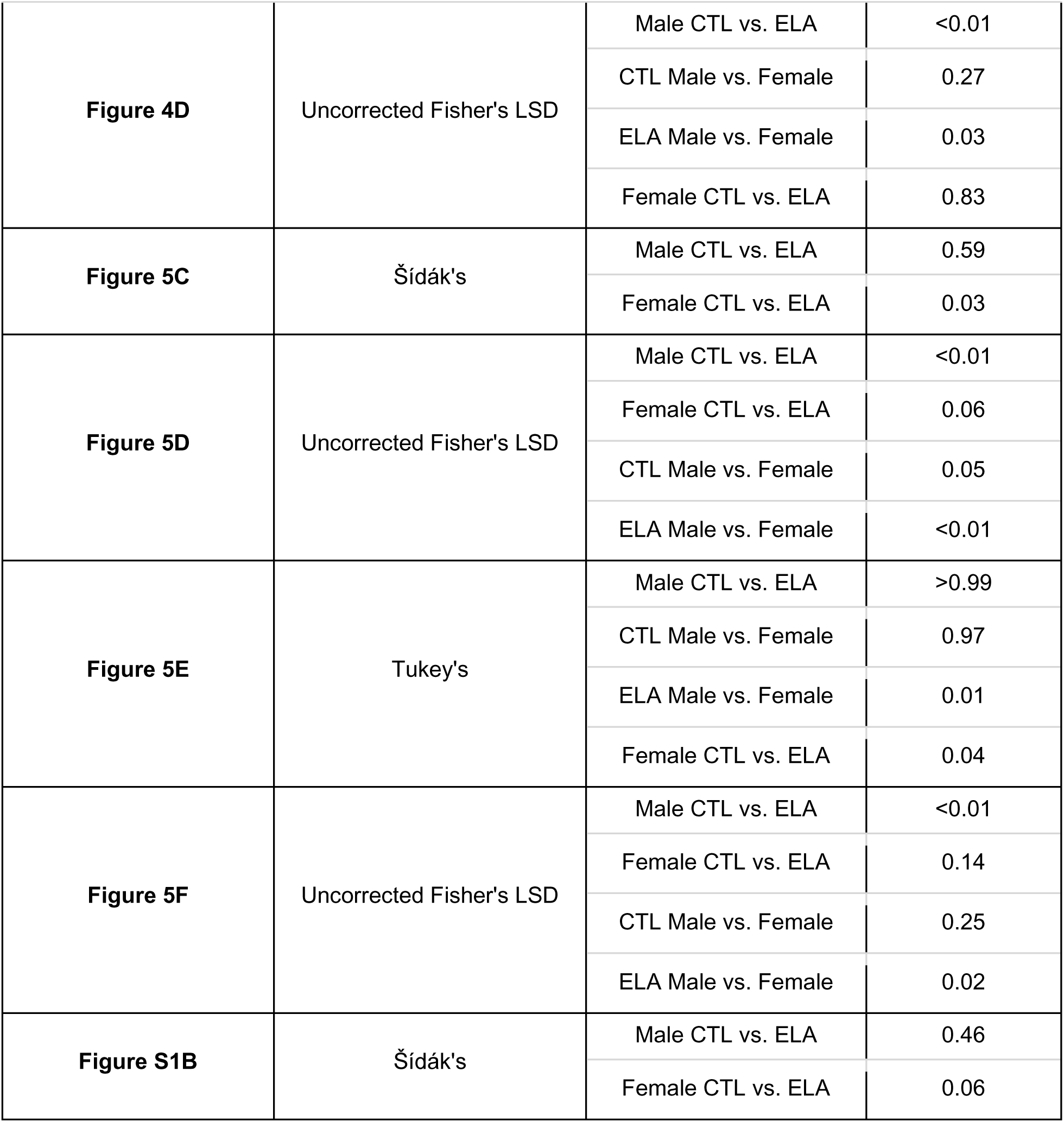
Multiple Comparison Tables.

| **Figure** | **Multiple Comparisons Table** | **Comparisons** | **Adjusted P Value** |
| --- | --- | --- | --- |
| **Figure 1B** | Uncorrected Fisher's LSD | Male CTL vs. ELA | <0.01 |
|  |  | CTL Male vs. Female | <0.01 |
|  |  | ELA Male vs. Female | 0.25 |
|  |  | Female CTL vs. ELA | <0.01 |
| **Figure 1C** | Uncorrected Fisher's LSD | Male CTL vs. ELA | 0.005 |
|  |  | Female CTL vs. ELA | 0.632 |
|  |  | CTL Male vs. Female | 0.213 |
|  |  | ELA Male vs. Female | <0.001 |
| **Figure 1D** | Uncorrected Fisher's LSD | Male CTL vs. ELA | 0.022 |
|  |  | CTL Male vs. Female | 0.1 |
|  |  | ELA Male vs. Female | <0.001 |
|  |  | Female CTL vs. ELA | 0.001 |
| **Figure 2B** | Šídák's | Male CTL VEH vs CNO | 0.02 |
|  |  | Female CTL VEH vs CNO | 0.83 |
| **Figure 2D** | Uncorrected Fisher's LSD | VEH Female CTL vs ELA | 0.29 |
|  |  | CNO Female CTL vs ELA | 0.03 |
| **Figure 3C** | Šídák's | CTL Male aCSF vs PTX | <0.001 |
|  |  | ELA Male aCSF vs PTX | <0.001 |
| **Figure 3G** | Šídák's | CTL Female aCSF vs PTX | <0.001 |
|  |  | ELA Female aCSF vs PTX | <0.001 |
| **Figure 3E** | Šídák's | CTL Male aCSF vs PTX | <0.001 |
|  |  | ELA Male aCSF vs PTX | 0.004 |
| **Figure 3H** | Šídák's | CTL Female aCSF vs PTX | <0.001 |
|  |  | ELA Female aCSF vs PTX | <0.001 |
| **Figure 4D** | Uncorrected Fisher's LSD | Male CTL vs. ELA | <0.01 |
|  |  | CTL Male vs. Female | 0.27 |
|  |  | ELA Male vs. Female | 0.03 |
|  |  | Female CTL vs. ELA | 0.83 |
| **Figure 5C** | Šídák's | Male CTL vs. ELA | 0.59 |
|  |  | Female CTL vs. ELA | 0.03 |
| **Figure 5D** | Uncorrected Fisher's LSD | Male CTL vs. ELA | <0.01 |
|  |  | Female CTL vs. ELA | 0.06 |
|  |  | CTL Male vs. Female | 0.05 |
|  |  | ELA Male vs. Female | <0.01 |
| **Figure 5E** | Tukey's | Male CTL vs. ELA | >0.99 |
|  |  | CTL Male vs. Female | 0.97 |
|  |  | ELA Male vs. Female | 0.01 |
|  |  | Female CTL vs. ELA | 0.04 |
| **Figure 5F** | Uncorrected Fisher's LSD | Male CTL vs. ELA | <0.01 |
|  |  | Female CTL vs. ELA | 0.14 |
|  |  | CTL Male vs. Female | 0.25 |
|  |  | ELA Male vs. Female | 0.02 |
| **Figure S1B** | Šídák's | Male CTL vs. ELA | 0.46 |
|  |  | Female CTL vs. ELA | 0.06 |

**Supplemental Table 3:** ANOVA Tables for Figure 5B.

| df = 11 for all | F values |  |  | P value |  |  |
| --- | --- | --- | --- | --- | --- | --- |
| Area | Sex | Rearing | Interaction | Sex | Rearing | Interaction |
| Frontal pole, cerebral cortex | 1.5046 | 0.0253 | 0.0013 | 0.246 | 0.876 | 0.972 |
| Primary motor area | 0.2591 | 1.3987 | 0.0724 | 0.621 | 0.262 | 0.793 |
| Secondary motor area | 1.4558 | 0.2112 | 0.8646 | 0.253 | 0.655 | 0.372 |
| Primary somatosensory area | 0.2258 | 0.8449 | 1.1143 | 0.644 | 0.378 | 0.314 |
| Supplemental somatosensory area | 0.0136 | 0.0835 | 4.3268 | 0.909 | 0.778 | 0.062 |
| Gustatory areas | 0.7096 | 0.0205 | 0.161 | 0.418 | 0.889 | 0.696 |
| Visceral area | 0.0172 | 0.473 | 0.0031 | 0.898 | 0.506 | 0.957 |
| Dorsal auditory area | 5.4343 | 11.8506 | 0.0988 | 0.04 | 0.006 | 0.759 |
| Primary auditory area | 1.2285 | 1.3894 | 1.0524 | 0.291 | 0.263 | 0.327 |
| Posterior auditory area | 0.1562 | 0.0274 | 0.1617 | 0.7 | 0.872 | 0.695 |
| Ventral auditory area | 0.1254 | 2.2449 | 0 | 0.73 | 0.162 | 0.998 |
| Anterolateral visual area | 4.9067 | 2.4006 | 0.9305 | 0.049 | 0.15 | 0.355 |
| Anteromedial visual area | 6.9037 | 2.9799 | 1.5307 | 0.024 | 0.112 | 0.242 |
| Lateral visual area | 5.389 | 2.8469 | 0.6597 | 0.04 | 0.12 | 0.434 |
| Primary visual area | 8.8056 | 7.3856 | 4.3112 | 0.013 | 0.02 | 0.062 |
| Posterolateral visual area | 5.1712 | 6.4261 | 0.0703 | 0.044 | 0.028 | 0.796 |
| Posteromedial visual area | 4.3588 | 4.5391 | 0.8527 | 0.061 | 0.057 | 0.376 |
| Anterior cingulate area | 3.839 | 1.5564 | 0.1401 | 0.076 | 0.238 | 0.715 |
| Prelimbic area | 1.9333 | 0.0054 | 0.0415 | 0.192 | 0.943 | 0.842 |
| Infralimbic area | 1.12 | 0.0004 | 1.6637 | 0.313 | 0.984 | 0.224 |
| Orbital area, lateral part | 1.0247 | 0.7006 | 0.199 | 0.333 | 0.42 | 0.664 |
| Orbital area, medial part | 3.1623 | 0.0883 | 0.5768 | 0.103 | 0.772 | 0.464 |
| Orbital area, ventrolateral part | 0.7356 | 0.0392 | 0.0001 | 0.409 | 0.847 | 0.991 |
| Agranular insular area, dorsal part | 0.1721 | 0.0044 | 1.0016 | 0.686 | 0.948 | 0.338 |
| Agranular insular area, posterior part | 2.9651 | 0.0116 | 1.3174 | 0.113 | 0.916 | 0.275 |
| Agranular insular area, ventral part | 0.0826 | 0.0142 | 2.3687 | 0.779 | 0.907 | 0.152 |
| Retrosplenial area | 20.8576 | 24.1938 | 3.0223 | 0.001 | 0 | 0.11 |
| Rostrolateral visual area | 3.3586 | 2.3914 | 0.0431 | 0.094 | 0.15 | 0.839 |
| Temporal association areas | 0.2928 | 0.5219 | 0.0013 | 0.599 | 0.485 | 0.972 |
| Perirhinal area | 1.9335 | 0.0571 | 0.0997 | 0.192 | 0.816 | 0.758 |
| Ectorhinal area | 0.0025 | 2.6946 | 0.083 | 0.961 | 0.129 | 0.779 |
| Olfactory areas | 0.2881 | 7.8988 | 3.4562 | 0.602 | 0.017 | 0.09 |
| Accessory olfactory bulb | 0.0473 | 1.1663 | 2.5289 | 0.832 | 0.303 | 0.14 |
| Anterior olfactory nucleus | 0.4856 | 1.2356 | 0.7711 | 0.5 | 0.29 | 0.399 |
| Taenia tecta | 0.4659 | 3.065 | 1.0047 | 0.509 | 0.108 | 0.338 |
| Dorsal peduncular area | 0.8584 | 1.0363 | 0.3313 | 0.374 | 0.331 | 0.577 |
| Piriform area | 3.0908 | 2.5585 | 1.8464 | 0.106 | 0.138 | 0.201 |
| Nucleus of the lateral olfactory tract | 0.116 | 0.0438 | 0.2245 | 0.74 | 0.838 | 0.645 |
| Cortical amygdalar area, anterior part | 0.004 | 0.1858 | 0.1299 | 0.951 | 0.675 | 0.725 |
| Cortical amygdalar area, posterior part | 0.473 | 1.523 | 0.1216 | 0.506 | 0.243 | 0.734 |
| Piriform-amygdalar area | 0.4598 | 1.7415 | 0.0275 | 0.512 | 0.214 | 0.871 |
| Postpiriform transition area | 1.594 | 4.257 | 2.8335 | 0.233 | 0.064 | 0.12 |
| Hippocampal formation | 0.67 | 0.5141 | 0.0163 | 0.43 | 0.488 | 0.901 |
| Field CA1 | 0.6297 | 5.2711 | 0.678 | 0.444 | 0.042 | 0.428 |
| Field CA2 | 0.0427 | 2.9538 | 1.5312 | 0.84 | 0.114 | 0.242 |
| Field CA3 | 0.0055 | 3.5865 | 1.551 | 0.942 | 0.085 | 0.239 |
| Dentate gyrus | 1.9501 | 1.4163 | 0.097 | 0.19 | 0.259 | 0.761 |
| Entorhinal area, lateral part | 3.3619 | 1.3072 | 0.6572 | 0.094 | 0.277 | 0.435 |
| Entorhinal area, medial part, dorsal zone | 0.004 | 0.2176 | 0.0369 | 0.951 | 0.65 | 0.851 |
| Parasubiculum | 1.4528 | 1.0061 | 0.7276 | 0.253 | 0.337 | 0.412 |
| Postsubiculum | 2.906 | 5.1989 | 2.0454 | 0.116 | 0.044 | 0.18 |
| Presubiculum | 0.701 | 6.0758 | 0.9257 | 0.42 | 0.031 | 0.357 |
| Subiculum | 1.4585 | 7.1208 | 2.2028 | 0.252 | 0.022 | 0.166 |
| Clastrum | 0.0176 | 0 | 0.1209 | 0.897 | 0.996 | 0.735 |
| Endopiriform nucleus | 7.8458 | 13.5283 | 1.517 | 0.017 | 0.004 | 0.244 |
| Posterior amygdalar nucleus | 0.2961 | 0.0067 | 0.1206 | 0.597 | 0.936 | 0.735 |
| Striatum | 0.0667 | 3.6612 | 0.4242 | 0.801 | 0.082 | 0.528 |
| Caudoputamen | 0.308 | 1.7991 | 0.5093 | 0.59 | 0.207 | 0.49 |
| Nucleus accumbens | 0.7107 | 6.3825 | 6.9421 | 0.417 | 0.028 | 0.023 |
| Olfactory tubercle | 0.2264 | 2.0741 | 2.0218 | 0.644 | 0.178 | 0.183 |
| Lateral septal complex | 0.0062 | 2.7488 | 0.1343 | 0.939 | 0.126 | 0.721 |
| Central amygdalar nucleus | 0.3407 | 3.2871 | 3.9402 | 0.571 | 0.097 | 0.073 |
| Medial amygdalar nucleus | 0.0859 | 0.5827 | 0.8567 | 0.775 | 0.461 | 0.375 |
| Pallidum | 0.455 | 1.1317 | 0.4409 | 0.514 | 0.31 | 0.52 |
| Globus pallidus, external segment | 0.3944 | 0.3555 | 0.0469 | 0.543 | 0.563 | 0.833 |
| Globus pallidus, internal segment | 0.1258 | 3.5662 | 0.2197 | 0.73 | 0.086 | 0.648 |
| Substantia innominata | 0.1144 | 17.8593 | 0.3159 | 0.742 | 0.001 | 0.585 |
| Magnocellular nucleus | 0.7262 | 1.465 | 0.029 | 0.412 | 0.252 | 0.868 |
| Medial septal nucleus | 0.0948 | 2.2114 | 0.112 | 0.764 | 0.165 | 0.744 |
| Diagonal band nucleus | 1.4319 | 0.0035 | 0.9576 | 0.257 | 0.954 | 0.349 |
| Triangular nucleus of septum | 2.3336 | 1.4979 | 0.2717 | 0.155 | 0.247 | 0.613 |
| Bed nuclei of the stria terminalis | 0.0788 | 2.639 | 1.1176 | 0.784 | 0.133 | 0.313 |
| Thalamus | 2.219 | 0.8193 | 0.0458 | 0.164 | 0.385 | 0.834 |
| Ventral group of the dorsal thalamus | 0.3732 | 0.2522 | 0.5928 | 0.554 | 0.625 | 0.458 |
| Subparafascicular nucleus | 0.1653 | 0.033 | 1.2753 | 0.692 | 0.859 | 0.283 |
| Subparafascicular area | 2.3651 | 0.2791 | 0.326 | 0.152 | 0.608 | 0.579 |
| Peripeduncular nucleus | 12.1556 | 2.0303 | 0.1774 | 0.005 | 0.182 | 0.682 |
| Geniculate group, dorsal thalamus | 0.2657 | 3.0176 | 0.0004 | 0.616 | 0.11 | 0.984 |
| Lateral group of the dorsal thalamus | 0.2642 | 0.3656 | 0.1507 | 0.617 | 0.558 | 0.705 |
| Anterior group of the dorsal thalamus | 0.2565 | 3.472 | 0.3026 | 0.623 | 0.089 | 0.593 |
| Medial group of the dorsal thalamus | 0.9554 | 0.2353 | 0.433 | 0.349 | 0.637 | 0.524 |
| Midline group of the dorsal thalamus | 0.9726 | 0.1802 | 1.7931 | 0.345 | 0.679 | 0.208 |
| Intralaminar nuclei of the dorsal thalamus | 1.6043 | 0.0795 | 0.8394 | 0.231 | 0.783 | 0.379 |
| Reticular nucleus of the thalamus | 0.0038 | 4.0715 | 0.6657 | 0.952 | 0.069 | 0.432 |
| Intergeniculate leaflet of the lateral geniculate complex | 3.9642 | 4.225 | 4.3418 | 0.072 | 0.064 | 0.061 |
| Ventral part of the lateral geniculate complex | 7.8283 | 6.6983 | 7.6344 | 0.017 | 0.025 | 0.018 |
| Subgeniculate nucleus | 3.3937 | 3.2641 | 3.8082 | 0.093 | 0.098 | 0.077 |
| Medial habenula | 0.8266 | 0.0214 | 1.7875 | 0.383 | 0.886 | 0.208 |
| Lateral habenula | 0.312 | 0.0002 | 0.1701 | 0.588 | 0.988 | 0.688 |
| Hypothalamus | 0.0419 | 1.5541 | 1.0628 | 0.842 | 0.238 | 0.325 |
| Periventricular zone | 0.3865 | 0.6141 | 0.0005 | 0.547 | 0.45 | 0.983 |
| Periventricular region | 0.8723 | 0.1726 | 0.444 | 0.37 | 0.686 | 0.519 |
| Anterior hypothalamic nucleus | 0.0022 | 0.1966 | 0.1293 | 0.964 | 0.666 | 0.726 |
| Mammillary body | 0.3243 | 0.9474 | 1.2456 | 0.58 | 0.351 | 0.288 |
| Medial preoptic nucleus | 0.0095 | 0.1666 | 0.0257 | 0.924 | 0.691 | 0.876 |
| Dorsal premammillary nucleus | 0.0004 | 0.0371 | 0.0931 | 0.985 | 0.851 | 0.766 |
| Ventral premammillary nucleus | 0.0499 | 4.6583 | 0.2557 | 0.827 | 0.054 | 0.623 |
| Paraventricular hypothalamic nucleus, descending division | 5.9314 | 0.026 | 0.1879 | 0.033 | 0.875 | 0.673 |
| Ventromedial hypothalamic nucleus | 0.0744 | 0.7946 | 0.6673 | 0.79 | 0.392 | 0.431 |
| Posterior hypothalamic nucleus | 9.659 | 0.0116 | 1.0793 | 0.01 | 0.916 | 0.321 |
| Hypothalamic lateral zone | 0.0863 | 4.0944 | 1.2698 | 0.774 | 0.068 | 0.284 |
| Zona incerta | 8.5572 | 4.0741 | 6.3245 | 0.014 | 0.069 | 0.029 |
| Median eminence | 0.0473 | 0.0002 | 1.451 | 0.832 | 0.989 | 0.254 |
| Midbrain | 6.3388 | 0.3981 | 0.4349 | 0.029 | 0.541 | 0.523 |
| Superior colliculus, sensory related | 12.8314 | 20.1473 | 15.7886 | 0.004 | 0.001 | 0.002 |
| Inferior colliculus | 0.0261 | 7.6096 | 0.5748 | 0.875 | 0.019 | 0.464 |
| Substantia nigra, reticular part | 0.0225 | 3.8425 | 0.02 | 0.884 | 0.076 | 0.89 |
| Ventral tegmental area | 3.3882 | 0.1994 | 0.5492 | 0.093 | 0.664 | 0.474 |
| Superior colliculus, motor related | 7.0314 | 6.2004 | 13.1079 | 0.023 | 0.03 | 0.004 |
| Periaqueductal gray | 0.6508 | 0.8526 | 0.26 | 0.437 | 0.376 | 0.62 |
| Precommissural nucleus | 0.8679 | 0.1714 | 0.4392 | 0.372 | 0.687 | 0.521 |
| Anterior pretectal nucleus | 11.3441 | 6.3047 | 8.2865 | 0.006 | 0.029 | 0.015 |
| Medial pretectal area | 3.5531 | 5.8071 | 4.4179 | 0.086 | 0.035 | 0.059 |
| Substantia nigra, compact part | 4.3916 | 6.113 | 4.5472 | 0.06 | 0.031 | 0.056 |
| Pedunclopontine nucleus | 1.793 | 1.7639 | 2.4251 | 0.208 | 0.211 | 0.148 |
| Interfascicular nucleus raphe | 3.2377 | 0.143 | 0.0824 | 0.099 | 0.712 | 0.779 |
| Interpeduncular nucleus | 1.9899 | 3.091 | 0.5291 | 0.186 | 0.106 | 0.482 |
| Rostral linear nucleus raphe | 64.9412 | 36.2273 | 26.0204 | 0 | 0 | 0 |
| Central linear nucleus raphe | 3.1103 | 1.1372 | 1.3354 | 0.106 | 0.309 | 0.272 |
| Dorsal nucleus raphe | 1.06 | 1.3254 | 0.0719 | 0.325 | 0.274 | 0.794 |

**Supplemental Table 4:** Abbreviations for Brain Regions in Figure 5B.

| **Abbreviation** | **Brain Region** |
| --- | --- |
| vStr | ventral striatum |
| NAc | nucleus accumbens |
| OT | olfactory tubercle |
| CeA | central amygdala |
| MeA | medial amygdala |
| SI | substantia innominata |
| MA | magnocellular nucleus |
| MS | medial septal nucleus |
| CA1 | field CA1 |
| CA2 | field CA2 |
| CA3 | field CA3 |
| DG | dentate gyrus |
| ENTl | entorhinal cortex |
| POST | postsubiculum |
| Pre | presubiculum |
| SUB | subiculum |
| EP | endopiriform cortex |
| SSs | somatosensory cortex, secondary |
| AUDd | auditory cortex, dorsal |
| VISal | anterolateral visual area |
| VISam | anteromedial visual area |
| VISl | lateral visual area |
| VISp | primary visual area |
| VISpl | posterolateral visual area |
| ACC | anterior cingulate cortex |
| Alp | Agranular insular area, posterior part |
| RSP | retrosplenial cortex |
| VISrl | Rostrolateral lateral visual area |
| ECT | ectorhinal cortex |
| OLF | olfactory areas |
| TT | taenia tecta |
| PIR | piriform cortex |
| NLOT | nucleus of the solitary tract |
| COA | cortical amygdalar area |
| PAA | piriform-amygdalar area |
| TR | postpiriform transition area |
| PP | peripeduncular cortex |
| GENd | geniculate group, dorsal thalamus |
| IGL | intergeniculate leaflet of the lateral geniculate complex |
| LGv | ventral part of the lateral geniculate complex |
| Subg | subgeniculate nucleus |
| LH | lateral habenula |
| LZ | hypothalamic lateral zone |
| ZI | zona incerta |
| MB | mammillary body |
| SCs | superior colliculus, sensory related |
| IC | inferior colliculus |
| SNr | substantia nigra, reticular part |
| VTA | ventral tegmental area |
| APN | anterior pretectal nucleus |
| MPT | medial pretectal area |
| SNC | substantia nigra, compact part |
| IF | interfascicular nucleus raphe |
| IPN | interpeduncular nucleus |
| RL | rostral linear nucleus raphe |
| CLI | central linear nucleus raphe |

## Supplemental Figures

**Supplemental Figure 1.**
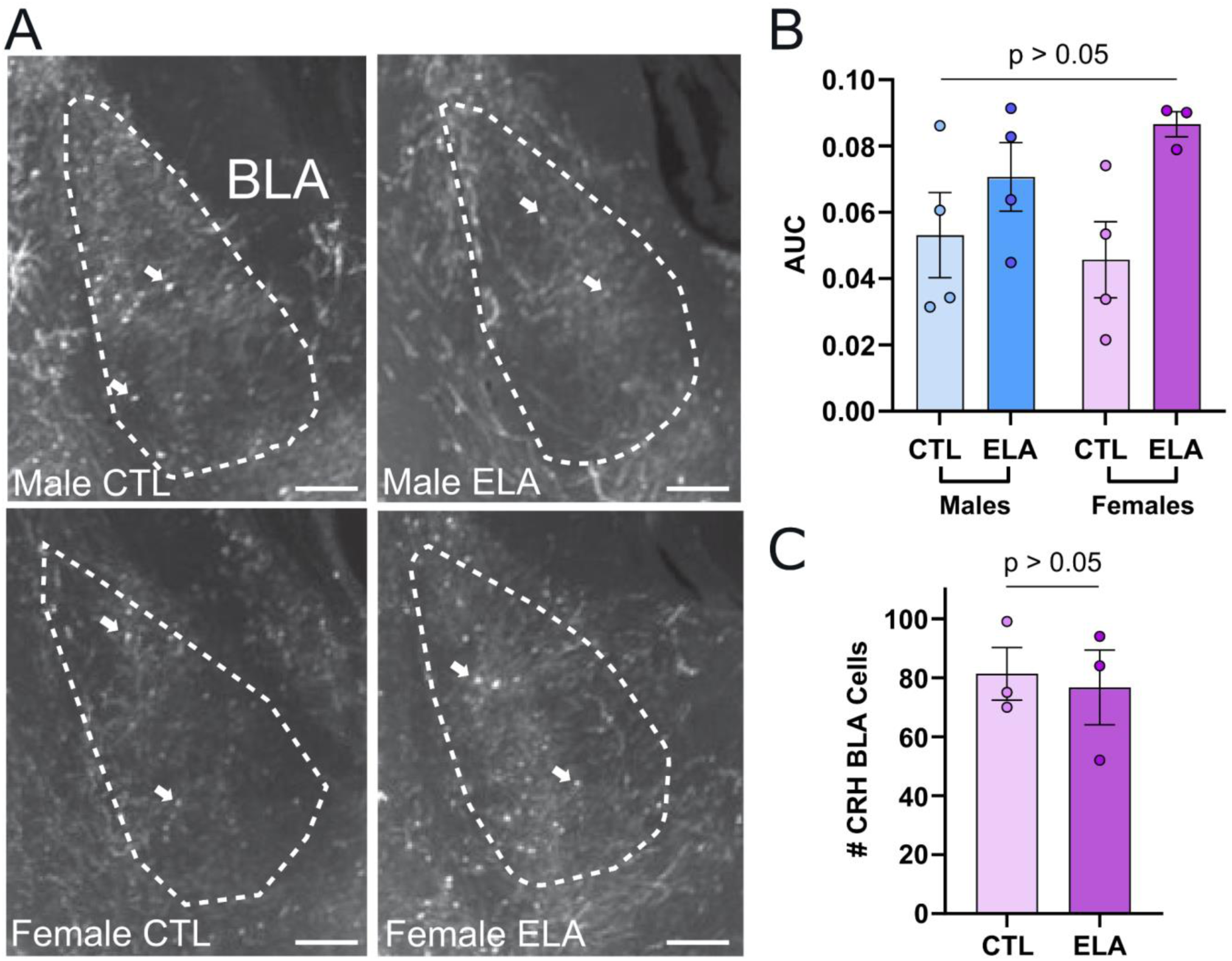
Viral labeling of CRH^+^ neurons in the BLA **(A)** Example 80 µm z-projections from raw images of posterior BLA in representative brains from each group. (**B**) Quantification of labeling as area under the curve (AUC). White arrows point to example labeling of CRH^+^ cell bodies. 2-way ANOVA (B: n, CTL male = 4, CTL female = 4, ELA male = 4, ELA female = 3). (**C**) Quantification of endogenous CRH cells in the BLA of CTL females and ELA females in 25 µm coronal sections, sampling from 4 sections per mouse. Unpaired t-test (C: n, CTL female = 3, ELA female = 3). CTL = control. ELA = early life adversity,

**Supplemental Figure 2:**
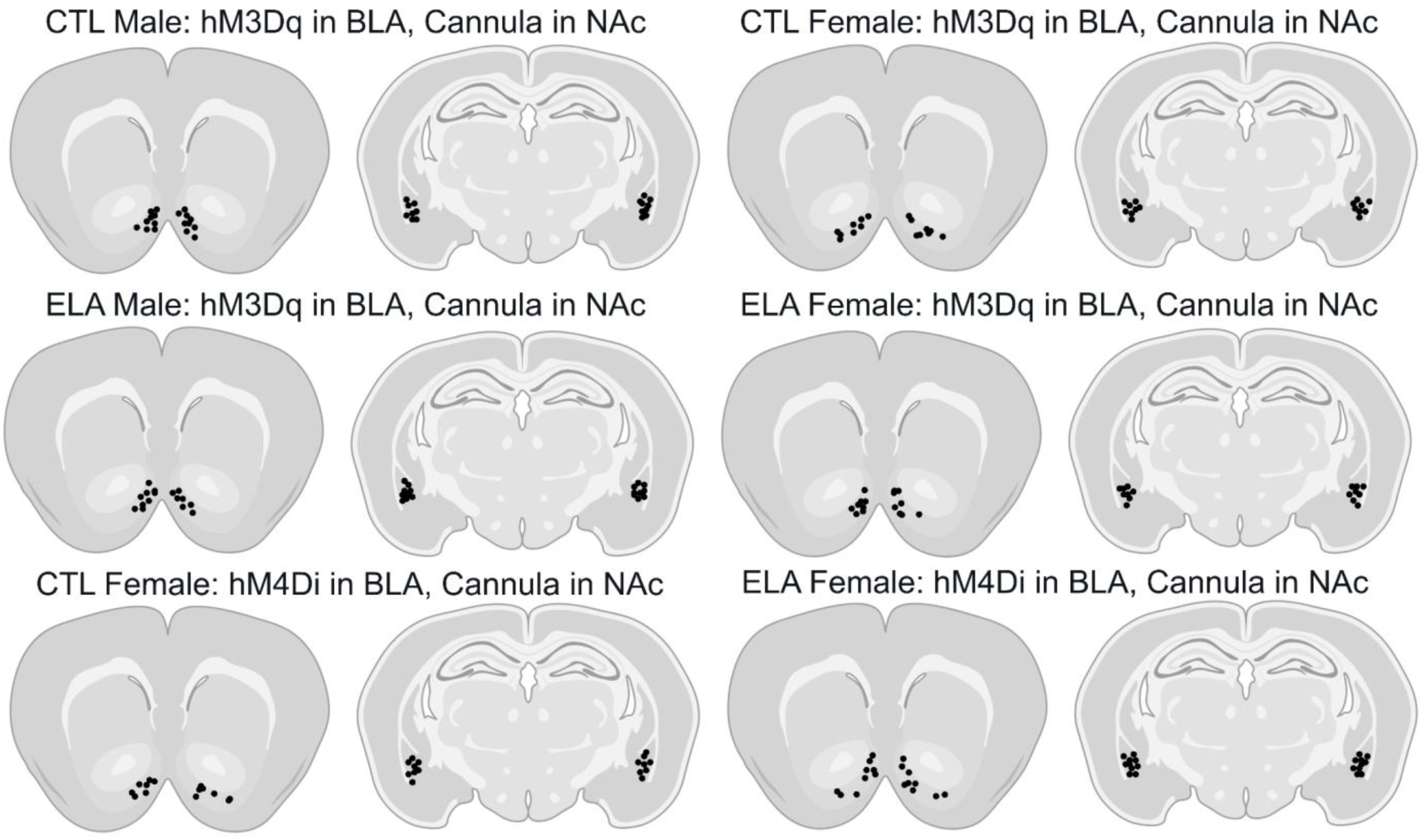
Viral injection and cannula fiber locations in the BLA and NAc. Schematic representations of bilateral viral injection sites and cannula locations of experiments in Figure 2.

